# CircPrime: a web-based platform for design of specific circular RNA primers

**DOI:** 10.1101/2022.12.20.521155

**Authors:** Fedor Sharko, Golam Rbbani, Prabhugouda Siriyappagouder, Joost A.M. Raeymaekers, Jorge Galindo-Villegas, Artem Nedoluzhko, Jorge M.O. Fernandes

## Abstract

**Background:** Circular RNAs (circRNAs) are covalently closed-loop RNAs with critical regulatory roles in cells. The tenth of thousands of circRNAs have been unveiled due to the recent advances in high throughput RNA sequencing technologies and bioinformatic tools development. At the same time, polymerase chain reaction (PCR) cross-validation for circRNAs predicted by bioinformatic tools remains an essential part of any circRNA study before publication.

**Results:** Here, we present the CircPrime web-based platform, providing a user-friendly solution for DNA primer design and thermocycling conditions for circRNA identification with routine PCR methods.

**Conclusions:** User-friendly CircPrime web platform (http://circprime.elgene.net/) works with outputs of the most popular bioinformatic predictors of circRNAs to design specific circular RNA primers. CircPrime works with circRNA coordinates and any reference genome from the National Center for Biotechnology Information database (NCBI).

## Background

In recent years, there is a marked increase in the number of circular RNA (circRNA)-related studies (Figure 1). CircRNAs have become a main focus of non-coding RNA biology research because they affect many genetic regulatory networks. These covalently closed-loop RNA molecules are an integral part of the cell regulome and interact with RNA-binding proteins. They can modulate microRNA expression and indirectly affect gene expression. In addition, some of them contain exon parts and can thus translate into proteins (Kristensen, et al., 2019).

**Figure 1.**
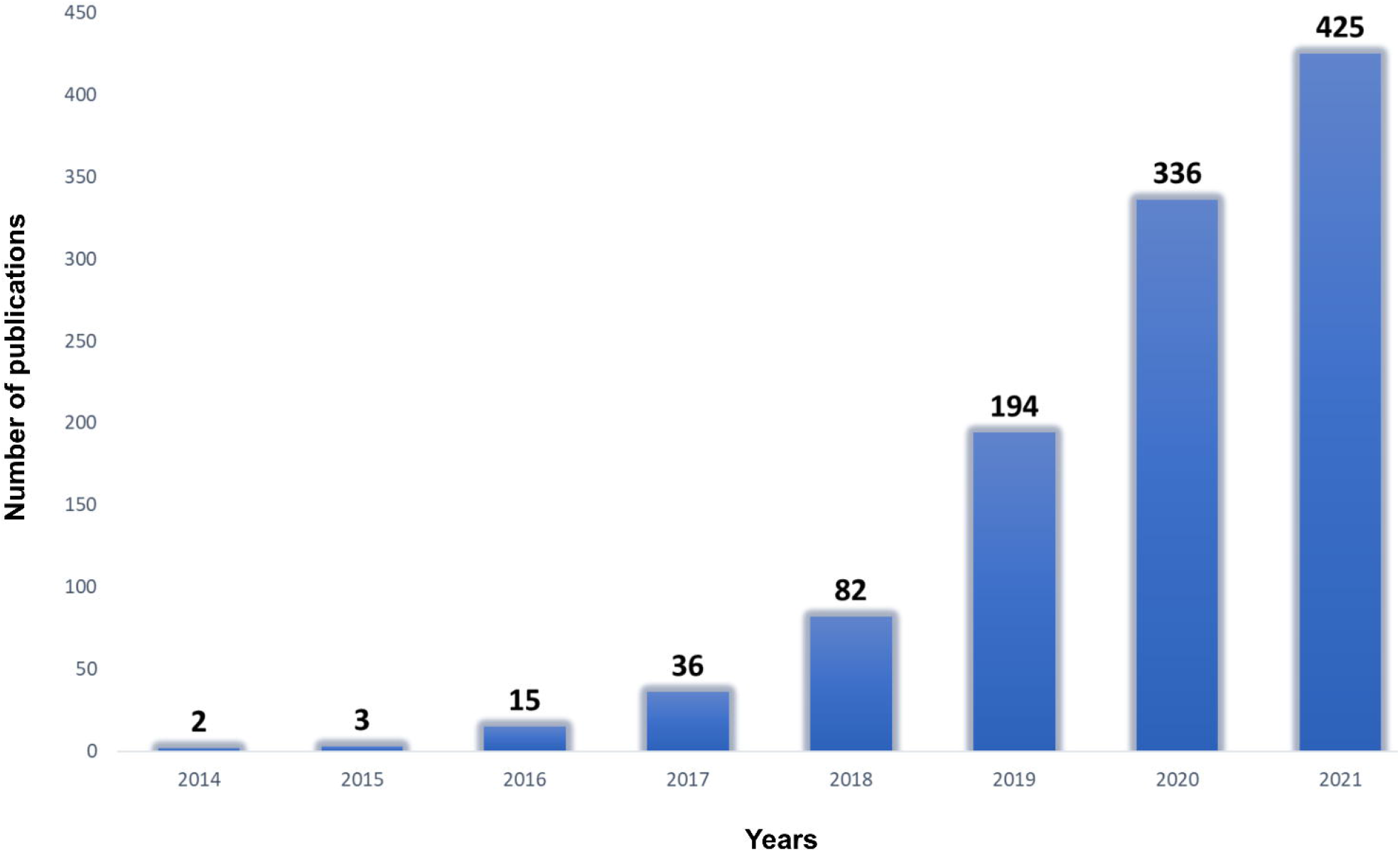
Exponential increase in the number of publications mentioning “circRNA” in their title. X-axis shows years from 2014 until 2021; Y-axis represents the number of publications. Source: Web of Science, accessed 6 October 2022.

Modern sequencing technologies make it now possible to identify hundreds of circRNAs that may be used as biomarkers and therapeutic targets in different applied researches (Nedoluzhko, et al., 2020; Rbbani, et al., 2021; Verduci, et al., 2021). However, *in silico* prediction of circRNAs leads to numerous false-positives (Hansen, et al., 2016), as well as inconsistencies among different bioinformatic pipelines (Nedoluzhko, et al., 2020). As a result, cross-checking and validation of circRNAs is an essential component of any circRNA study (Das, et al., 2022).

Reverse transcription PCR (RT-PCR) and quantitative PCR (qPCR) are considered the gold standard for identification of circRNA expression in cells (Das, et al., 2022). At the same time, primer design for the circRNAs validation differs from the design for the their linear host genes (Vromman, et al., 2022). To date, only a few tools have been published that allow the development of primers for validation of circRNAs. At the same time, they require additional software to be installed in different operating systems – CircPrimer (Zhong, et al., 2018), CircPrimer2.0 (Zhong and Feng, 2022) and circtools (Jakobi, et al., 2019), or work as a web tool with already known circRNAs of model organisms, namely human (Dudekula, et al., 2016) or novel circRNAs for limited number of animal species (Vromman, et al., 2022). Here, to overcome these previous constraints and facilitate the circRNA studies by presenting the user-friendly CircPrime web platform (http://circprime.elgene.net/), which works with outputs of the most popular bioinformatic predictors of circRNAs, such as CIRI2 (Gao, et al., 2018), KNIFE (Szabo, et al., 2015), CIRCexplorer2 (Zhang, et al., 2016), find_circ (Memczak, et al., 2013), circRNA_finder (Westholm, et al., 2014), DCC (Cheng, et al., 2016), mapsplice (Wang, et al., 2010) and common BED files. Importantly, CircPrime is also suitable for non-model organisms that have reference genome assemblies in the National Center for Biotechnology Information database (NCBI).

## Implementation

To date, there are several methods for PCR-based identification of different circRNAs types (Figure 2A). One of them is rolling circle amplification (RCA). This method avoids deep RNA sequencing and bioinformatic analysis, but is only capable of identifying a limited number of circRNA types (Boss and Arenz, 2020). The other most commonly used method assumes a longer workflow, which comprises circRNA enrichment, circRNA-library construction, deep sequencing, circRNA prediction, and finally RT-PCR/qPCR validation of bioinformatically predicted circRNAs (Shi, et al., 2022). PCR primers for this validation are designed to target the circRNA fragment overlapping a junction (back-splice) site of a specific circRNA (Figure 2A).

**Figure 2.**
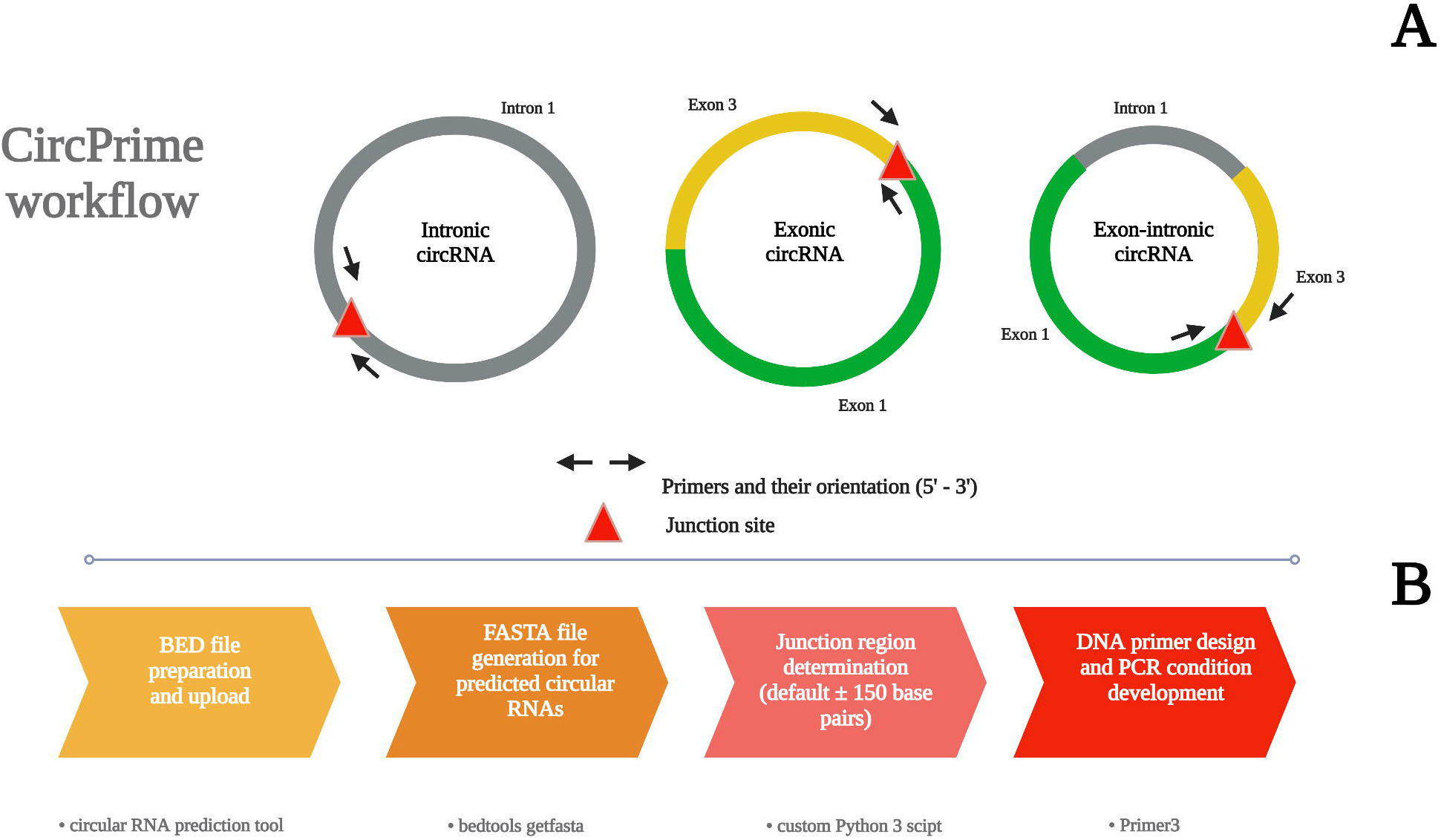
An overview of the CircPrime pipeline. (A) Types of possible variants for primer design for circular RNA validation. (B) The main steps of CircPrime pipeline and tools combined in it.

We developed CircPrime, as a streamlined pipeline in Python 3 and web platform, which makes use of output files from the most popular circRNAs *in silico* predictors. The CircPrime script implemented into the web platform currently contains the four main modules shown in Figure 2B and works under the parameters presented in Table 1.

**Table 1.** Required and default parameters for CircPrime usage.

| CircPrime parameter | Parameter description |
| --- | --- |
| CircRNA BED file | Required |
| Shift range left | Default 150 nucleotides |
| Shift range right | Default 150 nucleotides |
| Optimum length of a primer | Default 19 nucleotides |
| Minimum acceptable length of a primer | Default 15 nucleotides |
| Maximum acceptable length of a primer | Default 20 nucleotides |
| PCR product size | Default 150-200 nucleotides |
| Optimum melting temperature for a primer | in Celsius |
| Minimum acceptable melting temperature for a primer | in Celsius |
| Maximum acceptable melting temperature for a primer | in Celsius |
| Number of primer sets | Default 4 sets |

After the first step, which includes BED file uploading, CircPrime generates FASTA files using circRNA coordinates and reference genome from the NCBI. Then CircPrime extracts junction regions from the uploaded BED file and develops primer sets with the recommended melting temperature (Tm) for each circRNA in the list (up to 100) using Primer3 (Figure 2B) (Koressaar and Remm, 2007; Untergasser, et al., 2012). An example of the CircPrime output is presented as Supplementary Dataset 2.

## Results

The novel CircPrime web-based platform was evaluated to design primer sets for RT-PCR validation of circRNA expression in the muscle transcriptome of a teleost, the Nile tilapia (*Oreochromis niloticus*). Successfully, we showed that CircPrime significantly simplifies the primer design process for bioinformatically predicted circRNAs without the need to upload a reference genome of the organism studied.

In this study, we successfully applied circRNAs list predicted by CIRI2 (Gao, et al., 2018) and CircPrime web platform to design circRNA primer pairs and validate their RT-PCR efficiency using total RNA extract from the Nile tilapia skeletal muscle tissue (see details in Supplementary Material). We expect that this bioinformatic tool will play a relevant role on varied studies describing circRNAs expression and their possible functionality. CircPrime is applicable both on model and non-model organisms, including even those with a poor genome assembly and annotation.

## Conclusions

Herein, we present a Circprime web platform (http://circprime.elgene.net/) for PCR primer design and PCR conditions development for validation of circRNAs predicted based on RNA-sequencing data using different types of bioinformatics tools. We expect that this web tool will be convenient for users who intend to analyze the expression of circRNAs in animal and plant transcriptomes.

## Availability and requirements

Project name: CircPrime

Project home page: http://circprime.elgene.net/

Operating system(s): Platform independent

Programming language: Python 3.10

Other requirements: None

License: GNU GPL Version 3

## List of abbreviations

*PCR*: Polymerase chain reaction
*CircRNA*: Circular RNA
*RT-PCR*: Reverse transcription PCR
*qPCR*: Quantitative PCR
*NCBI*: National Center for Biotechnology Information database
RCA: Rolling circle amplification.

## Declarations

### Any restrictions to use by non-academics: no license needed

Ethics approval and consent to participate: This research was approved by the Nord University (Bodø, Norway) ethical committee. The experimental procedures involving animals were performed in accordance with the regulation and instructions of the Norwegian Animal Research Authority (FOTS ID 1042). All procedures involving animals were conducted according to the EU Directive 2010/63 on the use of animals for scientific purposes.

### Consent for publication

Not applicable

Availability of data and materials: The user-friendly CircPrime tool for circular RNA primer development is written in Python 3 and implemented on a web-based platform. It is freely available online at http://circprime.elgene.net/. The RNA-seq dataset generated and analysed during the current study is available in the GEO (NCBI) repository, under the accession number PRJNA826285

### Authors’ contributions

F.S. – wrote tool script and implemented it on a web-based platform, wrote and approved the final draft; G.R. – performed the experiments, prepared figures and/or tables, and approved the final draft; P.S. - performed the experiments, prepared figures and/or tables, and approved the final draft; J.R. – conceived and designed the experiments and approved the final draft; J.G. – conceived and designed the experiments and approved the final draft; A.N. – conceived and designed the experiments, performed the experiments, analyzed the data, prepared figures and/or tables, authored or reviewed drafts of the paper, and approved the final draft. J.M.O.F. – conceived and designed the experiments, authored or reviewed drafts of the paper, approved the final draft, administrated project and acquisited funding.

## Acknowledgements

Not applicable

