## Supplementary Material for "CircPrime: a web-based platform for design of specific circular RNA primers"

1. **Introduction**

This supplementary material contains the use case which supports our main manuscript regarding CircPrime: a web-based platform for the circular RNA primer design.

2. **Methods**
   2.1. **Ethical approval**

This research was approved by the Nord University (Bodø, Norway) ethical committee. The experimental procedures involving animals were performed in accordance with the regulation and instructions of the Norwegian Animal Research Authority (FOTS ID 1042). All procedures involving animals were conducted according to the EU Directive 2010/63 on the use of animals for scientific purposes.

2.2. **Sample collection**

Full-sibling male individuals of Nile tilapia (specimens: BM12; BM13; BM18; BM20) were randomly selected from the third generation of in-house domestication program (Podgorniak, et al., 2019) kept in a freshwater recirculating aquaculture system at Nord University's research station (Bodø, Norway). The system was maintained at temperature = 28 °C, pH = 7.5, photoperiod adjusted at 11:13 hours dark:light. All fish were fed ad libitum with 0.15–0.8 mm Amber Neptun pellets (Skretting, Norway). Fish were euthanatized with clove oil (Sigma Aldrich, USA) before sample collection using a 1:10 mix of 15 mL clove oil with 95% ethanol diluted in 10 L of system water. The fast (white) muscles were collected and frozen in liquid nitrogen by dissecting the left dorsal quadrant. Muscle samples were stored at -80°C until total RNA extraction.

2.3. **Total RNA extraction and quality control**

The frozen fast muscle samples were briefly homogenized in Lysing Matrix D, 2 mL tubes (MP Biomedicals, USA) with TRI reagent (Zymo Research, California, USA) at 6,500 rpm for 3 × 20s in a Precellys 24 homogenizer (Bertin Instruments, Montigny- le- Bretonneux, France). Total RNA was extracted from the homogenized tissue using the Direct-zol RNA kit (Zymo Research, California, USA) according to the manufacturers' instructions. RNA purity was assessed on a NanoDrop ND-1000 spectrophotometer (Thermofisher Scientific, USA) with the criteria of A260/280 ≥1.8 and A260/A230 ≥2.0. RNA concentration was evaluated using Qubit RNA Assay Kit with a Qubit 3.0 Fluorometer (Thermofisher Scientific, USA), and the RNA integrity number (RIN) was evaluated using TapeStation 2200 (Agilent Technologies, USA). Only RNA extracts with RIN higher than 7 were used for cDNA library preparation and subsequent sequencing.

2.4. **Circular RNA library preparation**

Approximately 500 ng of total RNA for each sample was used for circRNA library preparation. Total RNA was treated with RNase R (Lucigen, USA) for 10 min at 37 ˚C to digest all linear RNA and, the resulting RNA was then purified with Agencourt RNAClean XP beads (Beckman Coulter Inc., California, USA). Subsequently, NEBNext rRNA Depletion Kit v2 (Human/Mouse/Rat) with RNA Sample Purification Beads (NEB, Ipswich, USA) was used to ensure the complete removal of ribosomal RNAs (rRNAs). At the end of this procedure, RNA samples contain mainly circular RNA fragments. CircRNA multiplexed libraries were prepared using the NEBNext Ultra II RNA Library Prep Kit (NEB, Ipswich, USA). A unique barcode was tagged in each circRNA library and amplified with 16 PCR cycles. The quality and quantity of individual circRNA libraries were assessed using Agilent High Sensitivity D1000 ScreenTape assay in Agilent 2200 TapeStation (Agilent Technologies, USA). The multiplexed circRNA libraries were sequenced on Illumina NextSeq 500 platform (Illumina, San Diego, CA, USA) with the NextSeq 500 High Output Kit for 150 bp paired-end reads at the Nord University genomics facility (Bodø, Norway).

2.5. **Circular RNA prediction**

The DNA reads were filtered by quality (phred > 20) and library adapters were trimmed using fastp (v0.19.10) (Chen, et al., 2018), and further downstream analyses were performed with high-quality clean data. Reference genome (NCBI accession: GCF_001858045.2_O_niloticus_UMD_NMBU) and annotation file were downloaded from the National Center for Biotechnology Information (https://www.ncbi.nlm.nih.gov/). To identify circRNA, clean reads were mapped to Nile tilapia reference genome using BWA (v0.7.17) with these parameters: -T 19 -t 8, which uses a reference genome index for alignment (Li and Durbin, 2009). The resulting mapped reads from each sample were used as candidates for back-spliced junctions detection in CIRI2 v2.0.6 (with -T 4 parameter) algorithm (Gao, et al., 2018). Common circRNAs between sequenced libraries were subjected using online Venn diagram tool (<http://bioinformatics.psb.ugent.be/webtools/Venn/>). Host genes for common circRNAs were predicted using our previously published CircParser pipeline v0.7.1 (Nedoluzhko, et al., 2020).

2.6. **Primer design using CircPrime web-based platform**

We selected ten overrepresented circular RNAs from the CIRI2 output files which were overlapped between four circRNA datasets (Supplementary Dataset 1) and run CircPrime. We used the same Nile tilapia genome assembly version as previously; product size was expected as 201-600 bp. Shift range from the junction site was expected as 300 bp from each side of the fragment. Other parameters were used with the default options of CircPrime.

2.7. **PCR validation of circular RNAs using CircPrime-designed primers and gel-** **electrophoresis**

Primers and PCR conditions for circRNA validation were developed using CircPrime web platform. The same total RNA extracts used for the RNA library preparation were subjected to PCR to validate several high-expressed (with the highest number of junction reads) circRNAs. Complementary DNA (cDNA) was synthesized from the 200 ng total RNA using QuantiTect Reverse Transcription Kit (Qiagen, Germany). Divergent primers were expected to amplify the circRNAs fragment near the junction point. PCR amplification was performed with 40 cycles using AmpliTaq Gold DNA polymerase (Thermofisher Scientific, USA); subsequently, 2% agarose gel electrophoresis was used to visualize the PCR products.

3. **Results**

Four cDNA-libraries (BM12, BM13, BM18, BM20) were used for the deep sequencing (paired-end, 2 × 150 bp). The total number of reads generated for these four Nile tilapia DNA-libraries varied from 22,763,937 to 41,125,235. Post-filtering reads (after fastp quality filtration) were used for circRNA prediction using the CIRI2 algorithm (Gao, et al., 2018). Finally, from 241 to 386 circRNAs were identified (Figure S1A; Supplementary Dataset 1) and 28 of them were common between all libraries (Figure S1B; Supplementary Dataset 1).

We uploaded BED files for ten overrepresented, common for studied samples circular RNAs from the CIRI2 output files for primer design and PCR condition design to the CircPrime web tool. CircPrime was able to design primers sets for 9 from 10 circRNAs (Supplementary Dataset 2) and four of them (Table S1) were validated using PCR and were visualized by 2% (w/v) agarose gel electrophoresis (Figure S2).

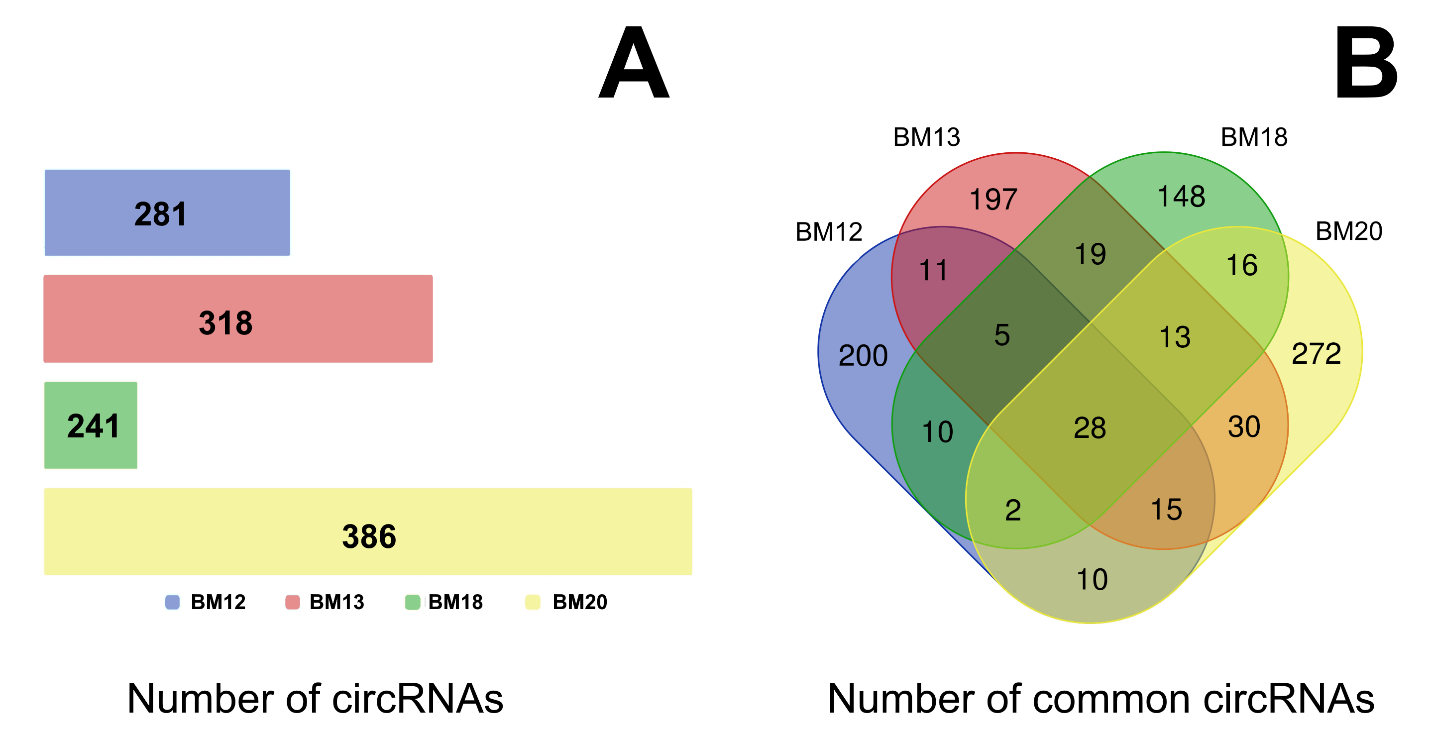

Figure S1. Circular RNAs that have been predicted by CIRI2 algorithm. (A) Number of circular RNAs that have been predicted for BM12, BM13, BM18, and BM20 cDNA libraries by CIRI2 algorithm. (B) Venn diagram with common circRNAs between BM12, BM13, BM18, BM20 cDNA libraries.

Table S1. CircRNAs and primer sets used for their PCR validation

| CircRNA / Host gene | Primer sequence 5’ – 3’ | Melting temperature, Tm |
| --- | --- | --- |
| circRNA1 / vasa gene for ATP-dependent RNA helicase DDX4 | Forward: CCCAGTACAACACGCCCAT | 58 °C |
|  | Reverse: GGCTGAAGCTTCTCACGCT | 58 °C |
| circRNA2 / titin (LOC100695257) | Forward: AAAGCCTTCACCTGCTCCC | 58 °C |
|  | Reverse: ACAAGATGCCAAGGTCACGA | 58 °C |
| circRNA3 / DDB1 and CUL4 associated factor 6 (dcaf6) | Forward: TGTCAAGTTGCTGGGACGG | 58 °C |
|  | Reverse: GCACTGGGTTTGAGCTCCT | 58 °C |
| circRNA4 / titin (LOC100702396) | Forward: TGAAGTGTGAGTGCGGTCC | 58 °C |
|  | Reverse: ACGGACTCCCCAGAGGTAC | 58 °C |

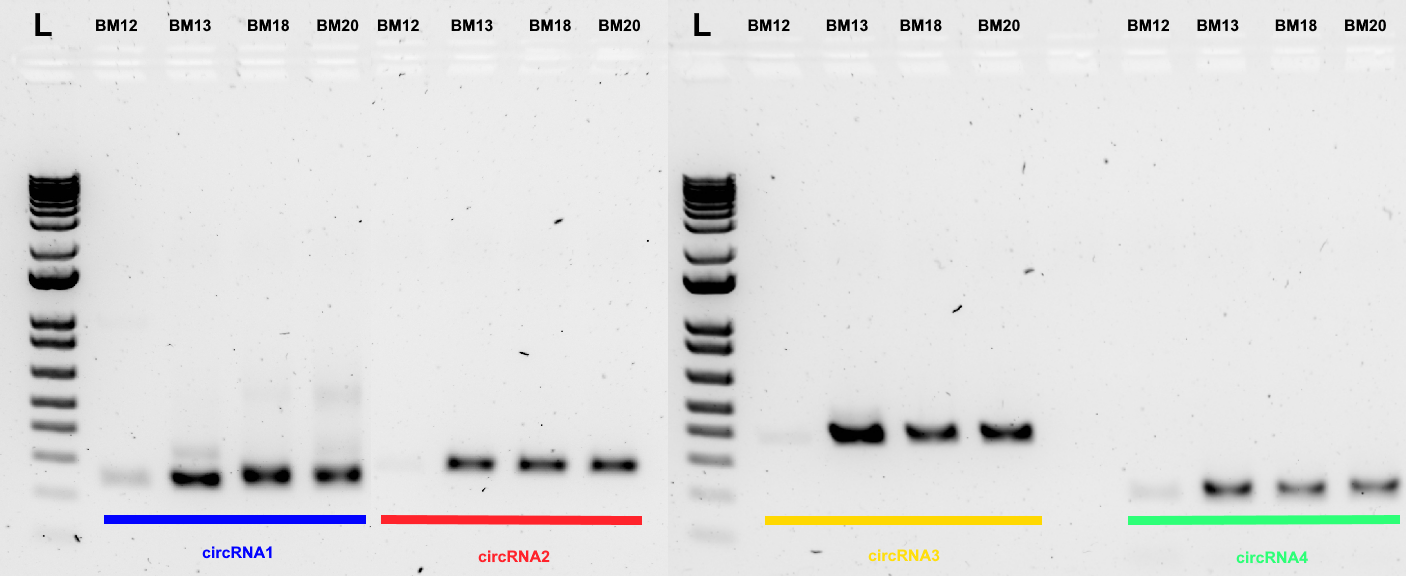

Figure S2. Agarose gel electrophoresis of PCR-positive circular RNAs. L in gel image represents 1 kb Plus DNA Ladder (Thermofisher Scientific, USA)

4. **Discussion**

Herein, we present a CircPrime web platform (<http://circprime.elgene.net/>) for PCR primer design and PCR conditions development for validation of circRNAs predicted based on RNA-sequencing data using different types of bioinformatics tools. We expect that this web tool will be convenient for users who intend to analyze the expression of circRNAs in animal and plant transcriptomes.

5. **References**

Chen, S.*, et al.* fastp: an ultra-fast all-in-one FASTQ preprocessor. *Bioinformatics* 2018;34(17):i884-i890.

Gao, Y., Zhang, J. and Zhao, F. Circular RNA identification based on multiple seed matching. *Brief Bioinform* 2018;19(5):803-810.

Li, H. and Durbin, R. Fast and accurate short read alignment with Burrows-Wheeler transform. *Bioinformatics* 2009;25(14):1754-1760.

Nedoluzhko, A.*, et al.* CircParser: a novel streamlined pipeline for circular RNA structure and host gene prediction in non-model organisms. *PeerJ* 2020;8:e8757.

Podgorniak, T.*, et al.* Differences in the fast muscle methylome provide insight into sex-specific epigenetic regulation of growth in Nile tilapia during early stages of domestication. *Epigenetics* 2019;14(8):818-836.

6. **Data availability**

The RNA-seq dataset generated and analysed during the current study is available in the GEO (NCBI) repository, under the accession number PRJNA826285
